# A coordinated regeneration–selection strategy enables genetic transformation and rapid flowering in apple × pear intergeneric hybrids

**DOI:** 10.64898/2026.04.09.717572

**Authors:** Furong Li, Miki Fujita, Mana Fujiwara, Yongbo Wang, Takuya Morimoto, Akihiro Itai

## Abstract

Wide hybridization between related species and genera provides valuable opportunities for broadening genetic diversity and introducing desirable traits. In the tribe Maleae (Rosaceae), *Malus* (apple) and *Pyrus* (pear) are phylogenetically closely related, and apple × pear hybrids represent promising materials such as for disease-resistance breeding. However, the effective utilization of such hybrids in breeding programs is constrained by long juvenile period. In this study, we established a tissue culture-based regeneration and genetic transformation platform for apple x pear hybrids. Key stages affecting adventitious shoot regeneration were optimized, and appropriate ranges of antibiotic selection pressure and bacterial elimination conditions were systematically evaluated. Regeneration capacity was predominantly genotype-dependent and became further restricted under *Agrobacterium* infection, necessitating precise balancing between regeneration competence and selection pressure. Using the highly competence line and the established transformation system, *MdFT1* gene was successfully introduced and over-expressed in intergeneric hybrids, resulting in transgenic plants exhibiting floral bud initiation approximately six months after infection under *in vitro* conditions. This study provides a practical and efficient regeneration–transformation framework for apple × pear hybrids and demonstrates its applicability for *FT-*mediated early flowering. The established system offers technical support for accelerated breeding strategies and facilitates the utilization of novel resources in genetic improvement of pome fruit.

## Introduction

Wide hybridization allows gene exchange across species or even genera and is an effective approach to broaden genetic diversity in crop plants^1^. It has been widely used to introduce agronomically important traits that are difficult to improve within a single species, including disease resistance, stress tolerance, and crop quality ^2–4^. Within the tribe Maleae of the family Rosaceae, *Malus* (apple) and *Pyrus* (pear) are phylogenetically related crops, which provides a biological basis for intergeneric hybridization and subsequent genetic introgression through backcrossing (Pasqualetto et al., 2023; Zhang et al., 2023). In recent years, increasing studies have reported the generation, molecular identification, and genetic characterization of intergeneric hybrids between *Malus* and *Pyrus*, supporting their feasibility as materials for intergeneric introgression breeding (Fischer et al., 2014; Morimoto et al., 2023). With respect to disease resistance, nonhost resistance to the respective scab pathogens (*Venturia spp*.) exists between *Pyrus* and *Malus* ^9^. Intergeneric resistance gene transfer from *Malus* to *Pyrus* have demonstrated that such resistance can be functionally exploited across closely related genera, thereby highlighting the strong application potential of intergeneric hybrids as practical breeding materials ^10^. Consistent with these findings, our previous work generated intergeneric hybrids between *Malus* and *Pyrus* and provided evidence that *Pyrus*-derived nonhost resistance to apple scab can be introgressed into an apple genetic background ^11^.

However, the stable incorporation of desirable traits derived from intergeneric hybridization into cultivated apple backgrounds generally requires repeated backcrossing and population selection (Migicovsky and Myles, 2017). In perennial fruit trees, the long juvenile phase markedly prolongs breeding cycles and represents a major bottleneck for the efficient utilization of intergeneric genetic resources. To shorten experimental and breeding timelines, genetic approaches aimed at inducing early flowering have been explored as tools for breeding acceleration ^13–15^

In apple and pear, early flowering has been achieved through manipulation of flowering-related gene expression or genome editing, with targeting of key regulators such as TERMINAL FLOWER 1 (TFL1) ^16^ and the MADS-box gene *BpMADS4* ^17^. In addition, FLOWERING LOCUS T (FT), a central component of florigen signaling, provides a rapid and visible phenotypic readout of transgene function, and its overexpression induces precocious flowering in both apple ^18^ and pear ^19^.

The effective application of such molecular tools in fruit cops critically depends on the availability of efficient and reproducible genetic transformation systems. Multiple regeneration and transformation systems have been developed independently for apple and pear (Chevreau et al., 2019; Xue et al., 2023; Maharjan et al., 2025). Meanwhile, these systems often exhibit pronounced genotype specificity ^24^ and are therefore difficult to directly transfer to intergeneric hybrid materials. In apple × pear intergeneric hybrids, the genetic complexity introduced by distant hybridization results in regeneration and transformation responses that cannot be reliably predicted from parental species. Therefore, it is necessary to establish an efficient and reproducible regeneration system for apple × pear intergeneric hybrids.

In this study, we established a tissue culture-based regeneration and genetic transformation platform for apple × pear intergeneric hybrids. Highly regenerable genotypes were first identified as suitable starting materials, followed by systematic optimization of key stages in adventitious shoot regeneration and culture conditions. Appropriate ranges of selection pressure and bacterial elimination were quantitatively evaluated to construct a reliable transformation pipeline. Using this platform, *MdFT1* gene was successfully introduced and expressed in intergeneric hybrids, resulting in transgenic plants exhibiting a flowering phenotype. These results together demonstrate that the established system enables stable gene delivery and functional gene expression in apple × pear intergeneric hybrids and provides a technical framework for accelerating genotype selection and trait evaluation through early flowering induction.

## Results

### Genotype-dependent regeneration capacity

Regeneration capacity differed markedly among six apple × pear intergeneric hybrids when cultured on four media M1–M4 (Fig.1). Although callus formation occurred in all genotypes, the magnitude of induction varied substantially. FjAD3-1, FjAD3-2, and FjOb3-1N consistently maintained high callus formation rates across media, whereas FjYaliB23-2 displayed strong medium dependence, with callus formation increasing from 36.67% on M1 to 91.11% on M4 (Fig. 2). Two-way ANOVA confirmed significant effects of genotype, medium, and their interaction (Supplementary Table S1).

**Fig. 1.**
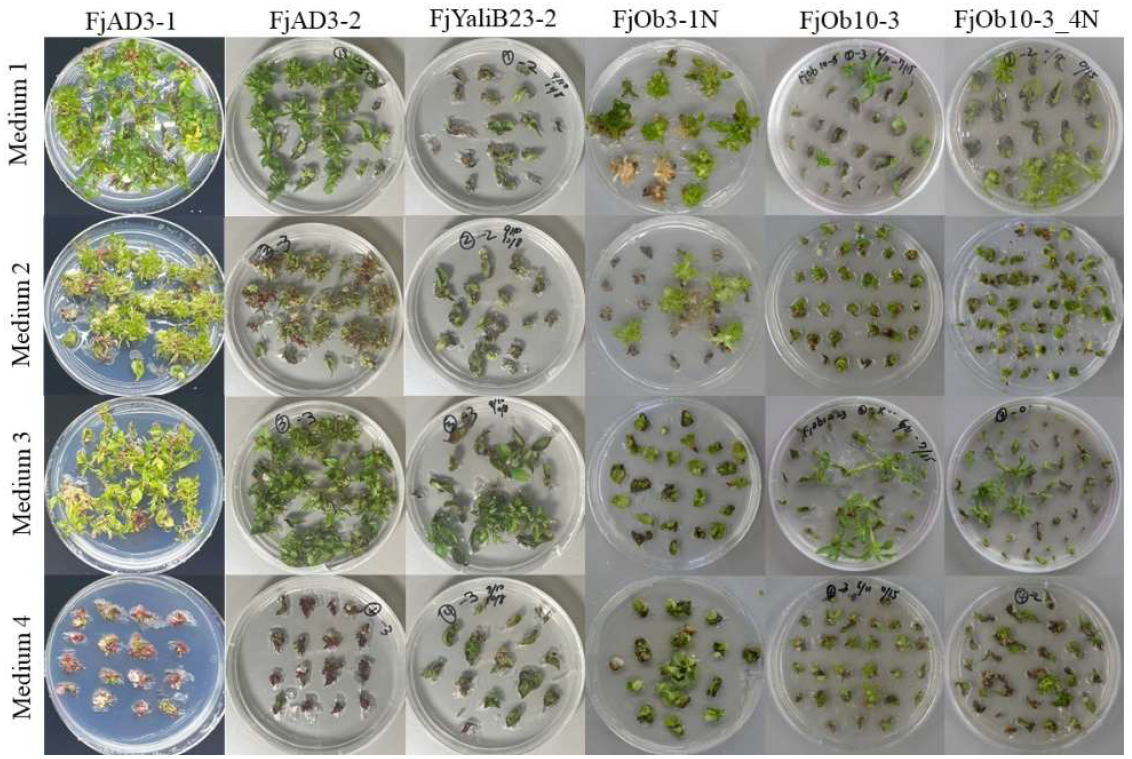
Regeneration responses of six intergeneric hybrid genotypes under four culture media. Explants from six apple × pear intergeneric hybrid genotypes were cultured on four regeneration media (M1–M4). Images were captured after 3 months of culture. Petri dishes were 90 mm in diameter.

**Fig. 2.**
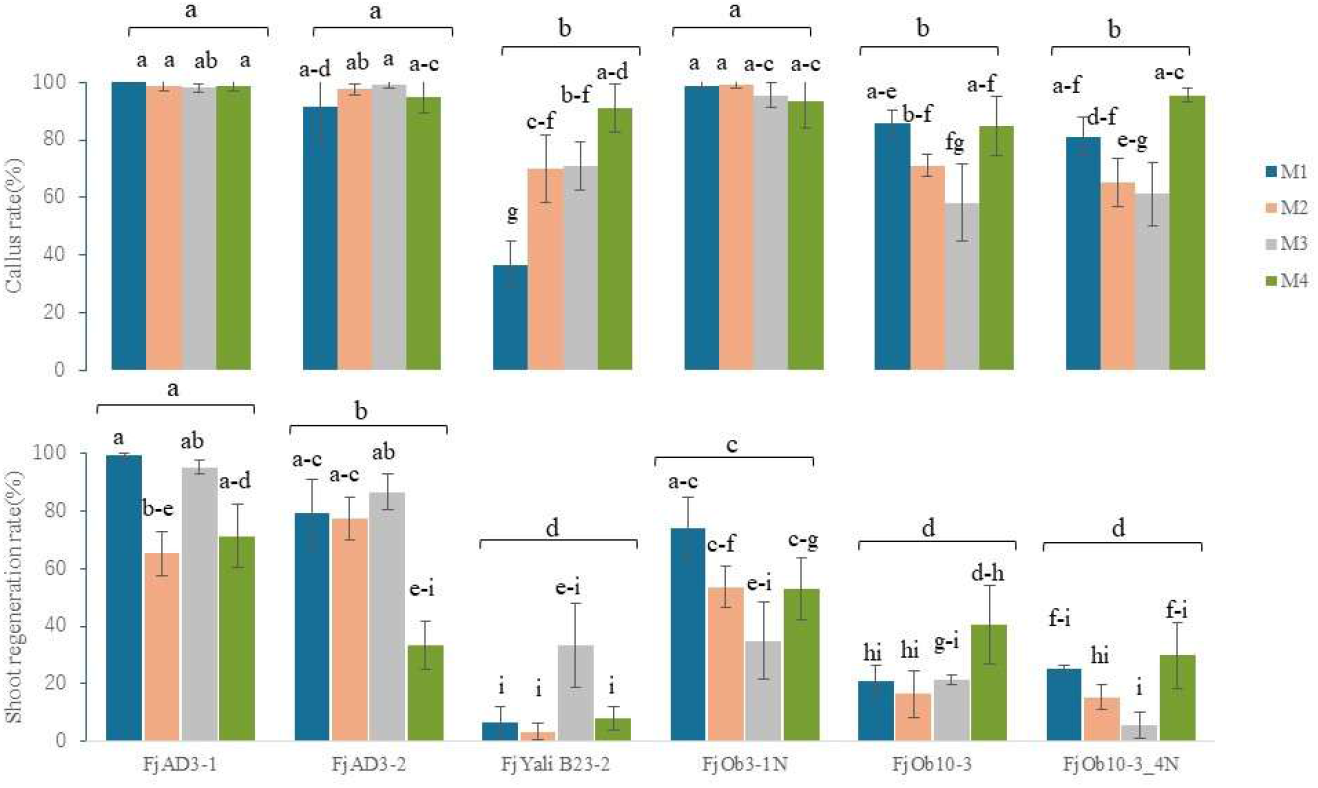
Interaction effects of genotype and culture medium on callus rate and regeneration rate. Data are presented as mean ± SD (n = 3, ≥30 leaves per replicate). Error bars indicate standard deviation. Different letters denote significant differences among variation (p < 0.05).

Differences among genotypes were more pronounced at the adventitious shoot regeneration stage. FjAD3-1 showed regeneration rates of 99.3% on M1 and 95.3% on M3, and maintained rates above 60% on M2 and M4. In contrast, FjYaliB23-2, FjOb10-3, and FjOb10-3_4N exhibited regeneration rates below 50% across all media, indicating persistently low regeneration capacity (Fig. 2). Two-way ANOVA further confirmed significant effects of genotype, medium, and their interaction on regeneration rate (Supplementary Table S2), with genotype contributing the largest statistical effect. Pooling regeneration data across the four media revealed the following ranking among genotypes: FjAD3-1 > FjAD3-2 > FjOb3-1N > FjOb10-3 = FjOb10-3_4N = FjYaliB23-2 (Fig. 2). Based on its consistently high and stable regeneration performance, FjAD3-1 was selected for subsequent transformation experiments.

FjAD3-1 was further evaluate with leaf developmental stage and media (Fig. 3). Compare with fully expanded leaves (Fig. 3a), the unexpanded leaves (Fig. 3b) showed higher regeneration (100%, Fig. 3d) on regeneration medium M1 and were therefore used in transformation experiments. Significant differences (p < 0.05) among media were observed for the time to first shoot emergence, shoot number per explant, and shoot length (Fig. 3c). M1 and M3 supported higher regeneration rates than M2 and M4. Notably, M1 promoted earlier shoot emergence (Fig. 3e), produced more shoots per explant (Fig. 3f), and enhanced shoot elongation (Fig. 3g). Overall, M1 provided the most balanced regeneration performance and was therefore selected for subsequent transformation experiments.

**Fig. 3.**
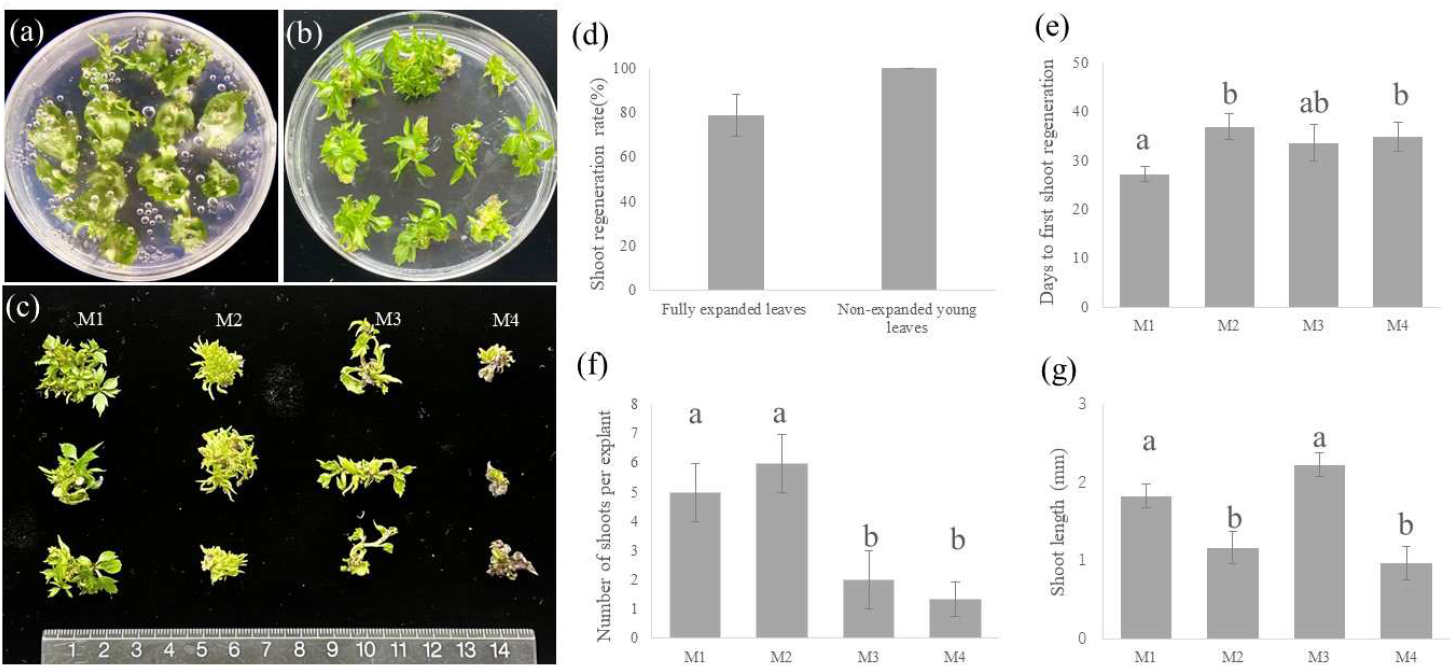
Effects of leaf developmental stage and regeneration medium on shoot regeneration in FjAD3-1. (a) Fully expanded leaves used as explants. (b) Unexpanded young leaves used as explants. (c) Regenerated shoots obtained on regeneration media M1–M4 after three months of culture. (d) Shoot regeneration rate of fully expanded and unexpanded young leaves with M1. (e) Time to first shoot emergence on media M1–M4. (f) Number of shoots per explant on media M1–M4. (g) Shoot length on media M1–M4. Data represent mean ± SD. Different letters indicate significant differences (P < 0.05).

### Optimization of antibiotic selection window

In the highly regenerative genotype FjAD3-1, the effect of Km concentration on adventitious shoot regeneration was evaluated under *Agrobacterium* free conditions to define an appropriate selection window for transformation. Regeneration rate declined sharply with increasing Km concentration (Fig.S1). While regeneration reached 100% in the absence of Km, it decreased to 47.9% at 5 mg L^−1^ and 18.6% at 10 mg L^−1^. At 15 mg L^−1^, regeneration was nearly abolished (2.6%), and no shoots were observed at 20 mg L^−1^ or higher (Fig. 4). These results indicated the high sensitivity of the selected apple × pear intergeneric hybrid line to Km selection pressure and identify 20 mg L^−1^ as a threshold concentration sufficient to fully suppress regeneration of non-transformed tissues. This threshold provided a reference for establishing effective selection conditions in subsequent transformation experiments.

**Fig. 4.**
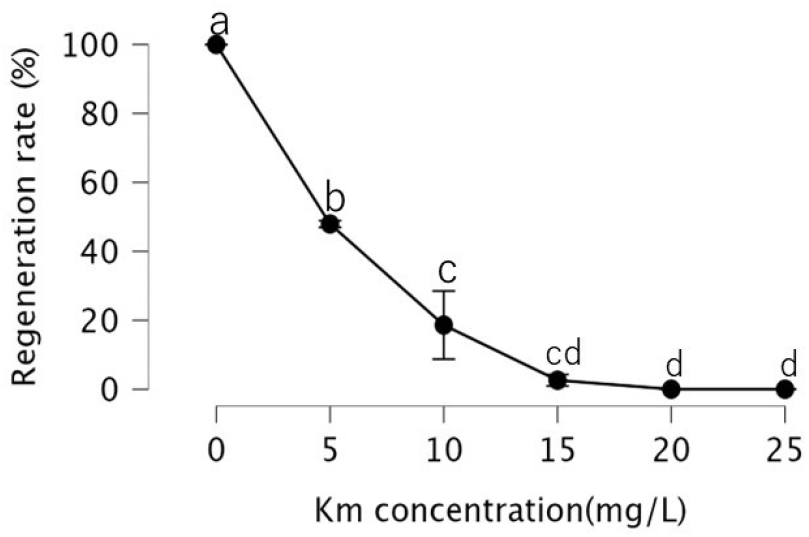
Effects of kanamycin concentrations on shoot regeneration. Data are presented as mean ± SD (n = 5). Different letters indicate significant differences among treatments (Tukey’s HSD test, p < 0.05).

The potential impact of the bactericidal antibiotic Me on regeneration was also assessed across concentrations ranging from 0 to 200 mg L^−1^. No significant differences in regeneration rate were detected among treatments (Fig. S2), indicating that Me did not interfere with shoot regeneration within the tested range. Accordingly, 50 mg L^−1^ Me was used for bacterial suppression during *Agrobacterium*-mediated transformation.

### Establishment of a transformation platform for apple × pear intergeneric hybrids

To establish a transformation system compatible with the regeneration framework of apple × pear intergeneric hybrids, *Agrobacterium*-mediated transformation experiments were conducted using the highly regenerative line, FjAD3-1, following determination of the Km sensitivity baseline. A binary vector carrying *nptII* as the selectable marker was used for transformation experiments. After co-cultivation, explants were transferred to selection media and monitored for adventitious shoot regeneration. Application of a high initial Km concentration (20 mg L^−1^) immediately after co-cultivation completely abolished regeneration (Table 1). Reducing the initial Km concentration to 15 or 10 mg L^−1^ permitted only minimal shoot formation, and regeneration rates remained markedly lower than those observed in non-infected explants at the same concentrations. These results suggest that *Agrobacterium* infection reduces regenerative capacity, effectively narrowing the selection window.

**Table 1.**
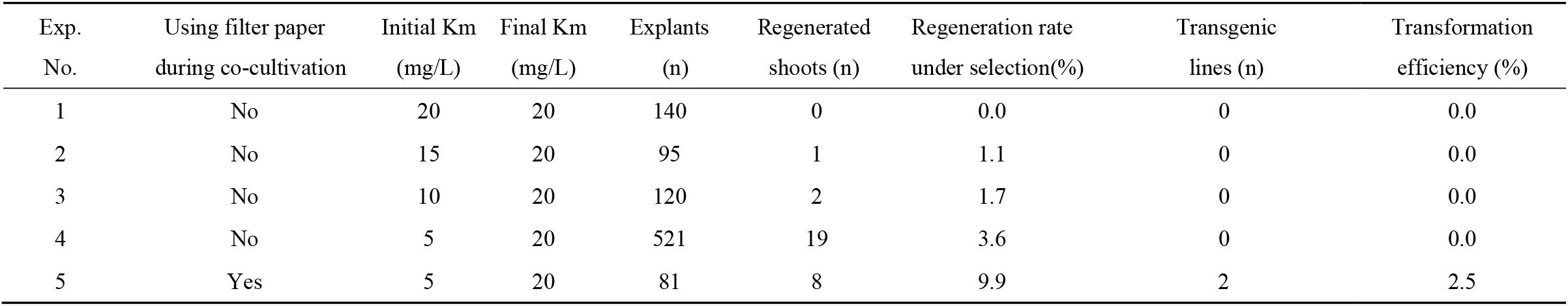
Effects of co-cultivation and selection conditions on regeneration and recovery of transgenic plants.

| Exp.<br>No. | Using filter paper<br>during co-cultivation | Initial Km<br>(mg/L) | Final Km<br>(mg/L) | Explants<br>(n) | Regenerated<br>shoots (n) | Regeneration rate<br>under selection(%) | Transgenic<br>lines (n) | Transformation<br>efficiency (%) |
| --- | --- | --- | --- | --- | --- | --- | --- | --- |
| 1 | No | 20 | 20 | 140 | 0 | 0.0 | 0 | 0.0 |
| 2 | No | 15 | 20 | 95 | 1 | 1.1 | 0 | 0.0 |
| 3 | No | 10 | 20 | 120 | 2 | 1.7 | 0 | 0.0 |
| 4 | No | 5 | 20 | 521 | 19 | 3.6 | 0 | 0.0 |
| 5 | Yes | 5 | 20 | 81 | 8 | 9.9 | 2 | 2.5 |

When the initial selection pressure was further reduced to 5 mg L^−1^, adventitious shoot regeneration became detectable (3.65%; Table 1). Under the same selection regime, incorporation of filter paper during the co-cultivation stage increased regeneration frequency to 9.9% (Table 1), corresponding to an approximately 2.7-fold increase without using filter paper. These findings suggest that maintaining regenerative competence under infection conditions requires both moderated early selection pressure and optimization of the co-cultivation microenvironment.

Under the optimized regime—filter paper-assisted co-cultivation, low initial selection (5 mg L^−1^), followed by stepwise increase to 20 mg L^−1^, stable Km-resistant shoots were recovered. Among 81 explants, 8 regenerated shoots were obtained, of which 2 were confirmed as PCR-positive transgenic lines (Fig. S3), corresponding to a final transformation efficiency of 2.5% (Table 1). These results define operational selection conditions compatible with regeneration under infection, providing a transformation platform for subsequent genetic manipulation.

### *MdFT1* overexpression induces early flowering in apple × pear intergeneric hybrids

Using the optimized transformation platform, transgenic lines overexpressing *MdFT1* under the control of the CaMV35S promoter were generated in the FjAD3-1 background. Following Km selection and subculture on shoot multiplication medium, regenerated shoots were subjected to molecular verification. PCR analysis confirmed the presence of the CaMV 35S-*MdFT1*-NosT expression cassette in regenerated shoots, whereas no amplification was detected in WT plants (Fig. 5a). RT–qPCR further revealed significantly elevated *MdFT1* transcript levels in transgenic lines relative to WT (Fig. 5b), indicating stable transgene expression.

**Fig. 5.**
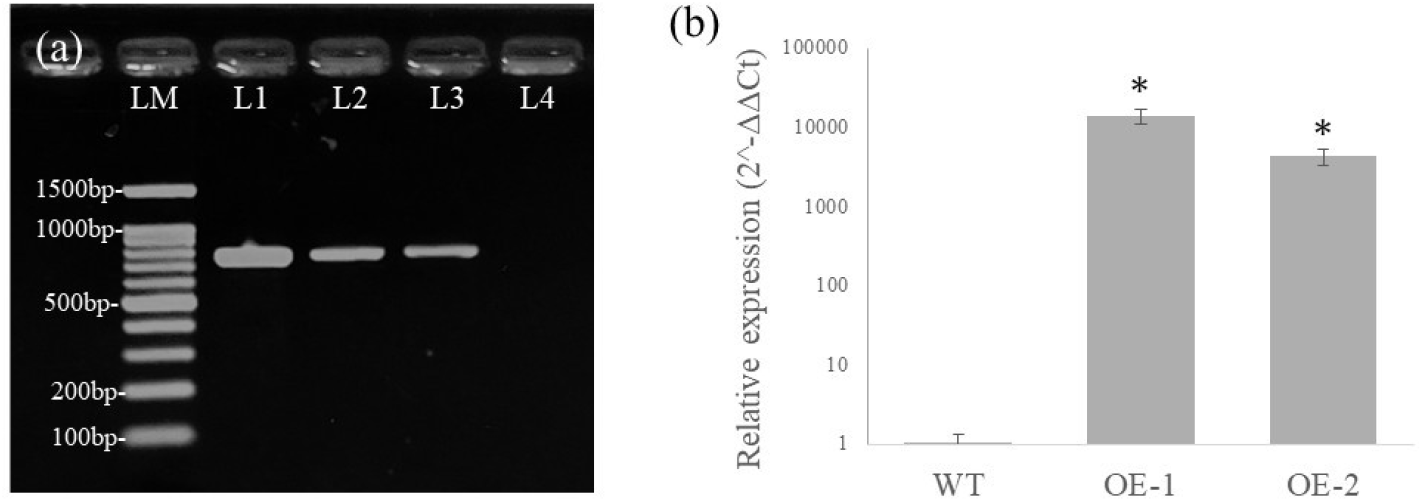
Molecular confirmation of *MdFT1* transgene integration and expression in apple × pear intergeneric hybrids. (a)PCR verification of the *MdFT1* transgene in transgenic lines. Lane M: 100 bp DNA ladder. Lane 1: plasmid containing the CaMV 35S-*MdFT1*-NosT gene used as a positive control. Lanes 2–3: transgenic lines showing the expected PCR amplification of the CaMV 35S-*MdFT1*-NosT gene (799bp). Lane 4: FjAD3-1 wild-type plant used as a negative control. (b) Relative *MdFT1* expression levels in transgenic lines determined by RT-qPCR. Expression levels were normalized to Actin and calculated using the 2^-ΔΔCt method. Error bars represent SD (n = 3). Asterisks indicate significant differences compared with WT (p < 0.05).

During continued maintenance on the shoot multiplication medium, *MdFT1*-transgenic plants initiated floral bud formation approximately six months after *Agrobacterium* infection (Fig. 6a). Under identical culture conditions, WT plants remained vegetative and did not exhibit floral induction. Floral buds were typically solitary (Fig. 6b), and differentiated anther structures were clearly visible within unopened buds (Fig. 6c). And floral buds subsequently developed and opened within culture vessels (Fig. 6d). Fully opened flowers exhibited morphologically normal floral organs including sepals, petals, anthers, and pistils.

**Fig. 6.**
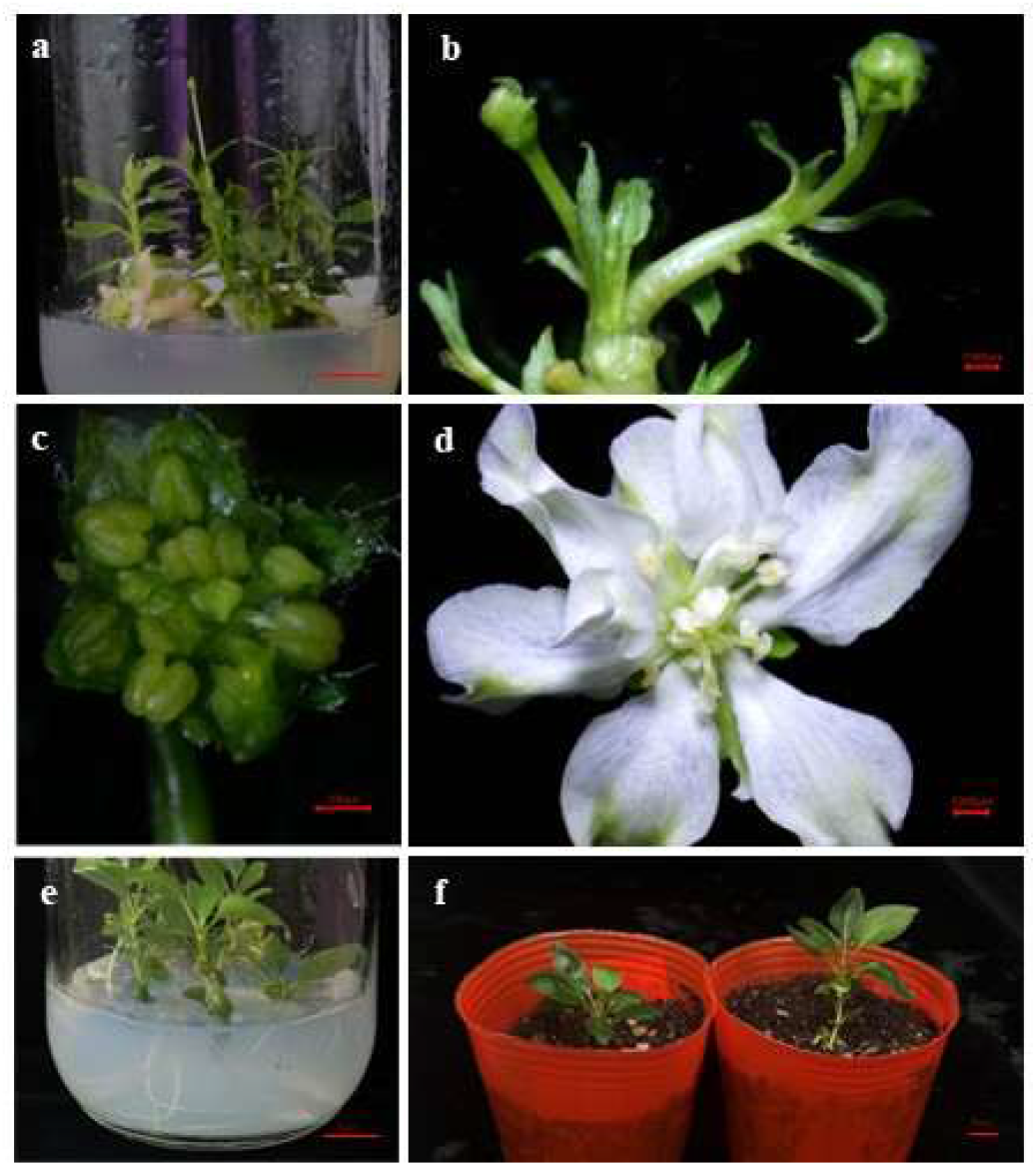
*MdFT1* overexpression induces early flowering and reproductive development in vitro in apple × pear intergeneric hybrids. (a) *MdFT1* transgenic plants flowering in vitro, 6 months after Agrobacterium infection. (b) Developing floral buds and flowers from *MdFT1* transgenic plants. (c) Stereomicroscopic observation of anthers after sepal removal. (d) Fully opened flower of transgenic plant. (e) Rooting of F *MdFT1* transgenic plantlets in vitro. (f) Acclimatized *MdFT1* transgenic plant grown in soil.

In addition, *MdFT1*-transgenic shoots retained rooting capacity under standard rooting conditions (Fig. 6e), indicating that accelerated reproductive transition did not compromise vegetative propagation ability. After transfer to soil, the transgenic plants were successfully acclimatized and continued normal growth (Fig. 6f). These results demonstrate that CaMV35S-driven *MdFT1* overexpression is sufficient to trigger early flowering in apple × pear intergeneric hybrids under standard *in vitro* propagation conditions while maintaining normal floral organ differentiation and rooting competence.

## Discussion

In plants, regeneration competence is shaped by genetic background, developmental state, and culture conditions (Long et al., 2022; Ochatt et al., 2025; Maharjan et al., 2025), leading to substantial variation in responses to *Agrobacterium*-mediated transformation. Such heterogeneity complicates the establishment of efficient and stable transformation systems. In woody perennials such as fruit trees, prolonged juvenile phases further delay the evaluation of reproductive traits and extend breeding cycles, increasing the importance of reliable transformation platforms. The challenge is intensified in intergeneric hybrids, which, despite offering valuable genetic diversity, often display unpredictable morphogenic responses that further constrain transformation efforts.

### Regeneration capacity is primarily genotype-dependent in apple × pear intergeneric hybrids

Genotype-dependent regeneration is a well-recognized phenomenon in woody perennials and has been widely reported in apple and pear systems (Dobránszki and Teixeira da Silva, 2010; Bell et al., 2012; Teixeira da Silva et al., 2019). Regeneration competence is strongly influenced by genetic background, often overriding the effects of medium composition or culture conditions. In intergeneric hybrids, where genomes from distinct genera are combined, such variability may become even more pronounced due to potential diversity in developmental regulatory networks. Consistent with this consensus, our results demonstrate that regeneration capacity in apple × pear intergeneric hybrids is predominantly determined by genotype. Although callus formation was observed across all six genotypes, adventitious shoot regeneration varied markedly among genetic backgrounds (Fig. 2). Statistical analysis revealed that the genotype effect was substantially stronger than the medium effect, indicating that intrinsic genetic factors play a dominant role in determining morphogenic competence in this intergeneric context. Notably, even hybrids sharing the same parental combination exhibited significant differences in regeneration performance (FjAD3-1 vs. FjAD3-2; FjOb3-1N vs. FjOb10-3), suggesting that recombination patterns in parental chromosomes rather than parental origin per se underlie regenerative divergence. By contrast, the tetraploid derivative FjOb10-3_4N did not display a significantly altered regeneration capacity relative to its diploid counterpart, suggesting that within the materials examined here, ploidy level did not substantially alter regeneration capacity.

Despite the predominant role of genotype, medium composition also exerted a significant influence on regeneration performance (Fig. 3). Significant genotype × medium interactions further indicate that the response to culture conditions varies among genetic backgrounds. Culture medium composition is widely regarded as a critical external factor influencing regeneration efficiency, primarily by modulating the expression of genotype-intrinsic regenerative potential through effects on cell division activity and organogenic processes ^29,31,32^. This concept has been well documented in woody fruit trees, including apple and pear, where culture conditions can partially release or suppress inherent regeneration competence in a genotype-dependent manner (Bommineni et al., 2001). Together, these findings suggest that culture conditions act as modulators of morphogenic expression within the regenerative potential defined by genetic background. Medium optimization improved regeneration efficiency within genotypic constraints but did not fully compensate for low-regenerating backgrounds under the conditions tested.

In addition to genotype and culture conditions, developmental stage of explant represents another critical determinant of regeneration outcomes ^34^. In the present study, unexpanded young leaves exhibited higher adventitious shoot regeneration rates and greater consistency among replicates compared with fully expanded leaves (3d), indicating improved regeneration stability. Younger explants generally exhibit higher regenerative potential than more mature tissues, with initial cell division preferentially occurring in younger regions near cambium and vascular tissues that are more responsive to hormonal cues ^25,35^.

Overall, regeneration capacity in apple × pear intergeneric hybrids is largely genotype dependent, with culture conditions and explant developmental stage exerting modulatory effects. Effective transformation platform in intergeneric materials therefore require coordinated consideration of genetic constitution, culture environment, and explant physiological state.

### Balancing antibiotic selection and regeneration capacity under Agrobacterium infection

Defining an appropriate selection window is a prerequisite for reliable recovery of transgenic events. In parental materials such as ‘Fuji’, Km at 10–25 mg L^−1^ has been reported to effectively suppress adventitious shoot regeneration (Hyung et al., 2003; Arcos et al., 2020; Cárdenas and Stange Klein, 2025). In contrast, the apple × pear intergeneric hybrids examined here exhibited pronounced regeneration inhibition at 5 mg L^−1^ under non-infected conditions (Fig. 4), indicating a lower tolerance threshold to Km in this genetic background. Complete suppression of regeneration occurred at 20 mg L^−1^ (Fig. 4), thereby defining the upper reference boundary for selection intensity in this material.

Under Agrobacterium infection conditions, regeneration was further constrained. Application of 20 mg L^−1^ Km resulted in the absence of shoot regeneration, while 5–15 mg L^−1^ permitted only minimal regeneration frequencies (Table 1). In contrast, Me had no detectable effects on regeneration within the tested range (Fig. S2), consistent with previous reports in apple transformation studies (Li et al., 2011; Morinaka et al., 2022). Collectively, these findings indicate that *Agrobacterium* infection substantially narrows the effective selection window, representing a key limitation for transformation efficiency in complex intergeneric materials. Efficient recovery of transgenic events therefore requires balancing selection pressure with the intrinsic regeneration capacity of the target material ^40,41^.

The use of filter paper during co-cultivation, as reported, is thought to moderate excess bacterial proliferation and reduce tissue stress, thereby facilitating subsequent regeneration ^42,43^. Under identical selection conditions, filter paper-assisted co-cultivation increased regeneration frequency under selection approximately 2.7-fold (Table 1). Although overall transformation efficiency remained limited, this combined optimization increased the probability of recovering transgenic events.

Despite these improvements, overall transformation recovery remained low, highlighting the inherent recalcitrance of apple × pear intergeneric hybrids. Further optimization will therefore be required, including refinement of selection timing ^44^, adjustment of pre-culture conditions (Li et al., 2011; Lin et al., 2025), improvement of infection parameters ^46,23^, and enhancement of regeneration competence through morphogenic regulators^47^, to increase the proportion of cells competent for stable transformation. Together, these findings underscore that transformation recovery in complex intergeneric hybrids requires coordinated adjustment of regenerative capacity and selection strength under infection.

### FT-mediated acceleration of reproductive transition in intergeneric hybrids

FT functions as a central floral integrator and has been widely reported to induce early flowering in diverse woody species (Tränkner et al., 2010; Zhang et al., 2010; Tomes et al., 2023). In the present study, CaMV35-driven *MdFT1* overexpression effectively triggered reproductive transition in apple × pear intergeneric hybrids approximately six months after *Agrobacterium* infection under *in vitro* conditions (Fig. 6). Notably, *MdFT1-*induced early flowering was not accompanied by obvious morphological abnormalities, although the functionality as reproductive organs were not systematically evaluated in the present study. Whether the *in vitro* flowering phenotype remains stable under greenhouse or field conditions warrants further investigation.

Intergeneric hybridization between apple and pear provides valuable genetic resources for breeding, particularly for the introgression of disease resistance and other complex traits. Nonhost resistance to the apple scab pathogen (*Venturia inaequalis*), for example, has been reported in pears ^9^. Importantly, our previous study^11^ demonstrated that such pear-derived nonhost resistance can be transmitted to apple × pear hybrids, providing a practical basis for its utilization in apple breeding through intergeneric hybridization. However, practical utilization of such materials is often constrained by long juvenile phases and the lack of reliable regeneration-transformation systems, which together slow generation turnover and hinder precise genetic manipulation. In this study, coupling a regeneration-optimized transformation platform with *MdFT1*-mediated early flowering provides a practical approach to shorten the juvenile phase. Although early flowering does not replace field evaluation, reducing generation time is particularly valuable for breeding programs requiring repeated backcrossing or trait introgression.

In summary, this work highlights that intergeneric materials can support stable genetic manipulation when regeneration competence and selection dynamics are properly coordinated. The platform described here provides a practical foundation for genome editing in apple × pear hybrids and for targeted trait introduction from both genera.

## Materials and Methods

### Plant materials

Intergeneric hybrid lines between apple (*Malus* × *domestica*) and pears (*Pyrus* spp.) used in this study were originally generated by our previous study using controlled crosses followed by seeding development *in vitro* (Morimoto et al., 2020; 2024). The hybrid lines included FjOb3-1N and FjOb10-3, derived from a crosse between apple ‘Fuji’ and pear interspecific hybrid (*P. pyrifolia* × *P. communis*) ‘Oharabeni’; FjAD3-1 and FjAD3-2, derived from a crosse between ‘Fuji’ and European pear (*P. communis*) ‘Alexandrine Douillard’; and FjYali23-2, derived from a cross between ‘Fuji’ and Chinese white pear (*P. bretschneideri*) ‘Yali’, as previously described. FjOb10-3 was further used to induce tetraploid individuals resulting in FjOb10-3_4N. All lines were maintained *in vitro* on shoot multiplication medium (MS medium supplemented with 1 mg L^−1^ 6-BA and 0.1 mg L^−1^ IBA) and subcultured every 4 weeks under sterile conditions.

### Optimization of shoot regeneration assay

We selected four regeneration media, M1–M4 (Table 2), based on previously reported leaf explant regeneration systems for apple and pear (Maximova et al., 1998; Matsuda et al., 2005; Li et al., 2018; Kawato et al., 2023). To compare regeneration capacity, six apple × pear intergeneric hybrids lines were cultured on the four regeneration media formulations. Young leaves including unexpanded and fully expanded leaves were excised from the apical region of shoots that had been cultured *in vitro* for four weeks. Leaves were placed flat on regeneration media. Cultures were maintained at 25 °C, with the first two weeks under dark conditions, followed by two weeks under low-light conditions, and subsequently transferred to a 16 h light / 8 h dark photoperiod. Culture media were renewed monthly. To evaluate the effect of leaf developmental stage, FjAD3-1 were further evaluated for regeneration capacity between unexpanded and fully expanded young leaves.

**Table 2.**
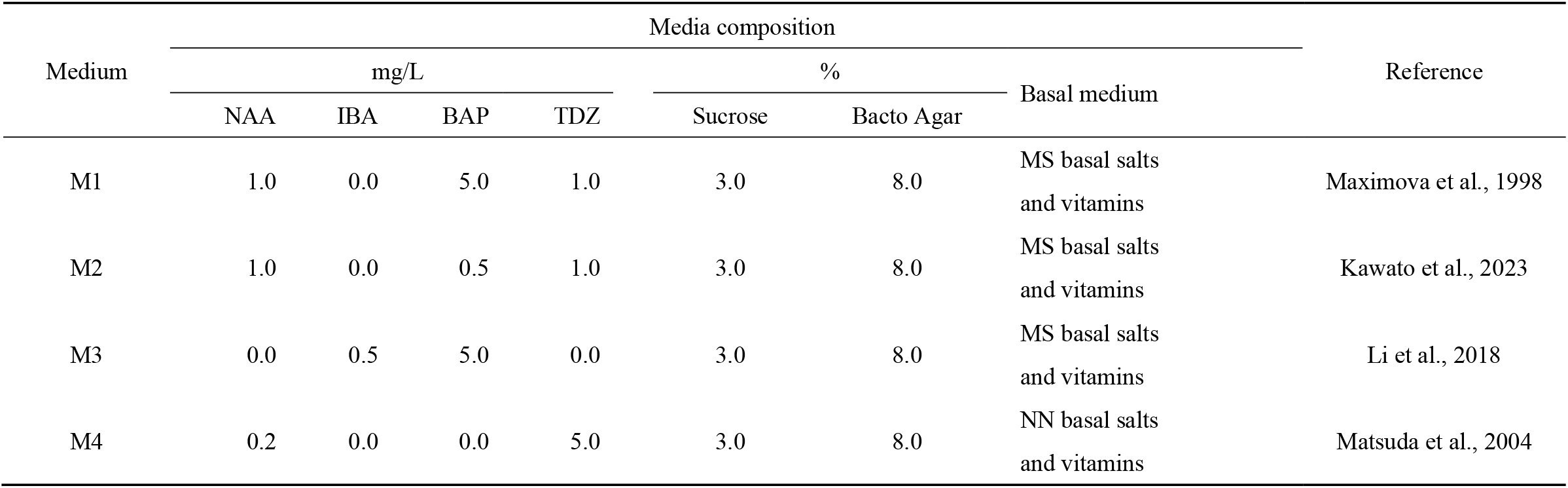
Composition of the regeneration media.

Callus formation rate (%) was calculated as the number of explants forming callus divided by the total number of explants × 100. Adventitious shoot regeneration rate (%) was calculated as the number of explants producing adventitious shoots (≥ 2 mm in length) divided by the total number of explants ×100. The number of shoots per explant was defined as clearly distinguishable adventitious shoots (≥ 2 mm in length). Shoot length was measured from the base of the shoot to the apex. Time to first shoot emergence was recorded as the number of days from the initiation of explant culture to the appearance of the first visible adventitious shoot.

Each treatment consisted of three biological replicates, with at least 30 explants per replicate. Regeneration parameters were recorded after three months of culture. Data were analyzed using biological replicates as statistical units and are presented as mean ± standard deviation (SD).

### Determination of selection and bacterial elimination conditions

To determine appropriate antibiotic conditions for selection and bacterial elimination, kanamycin (Km) and meropenem (Me) were added to the regeneration medium M1 at different concentrations. Km was tested at 0, 5, 10, 15, 20, and 25 mg L^−1^, and Me at 0, 50, 100, 150, and 200 mg L^−1^. All other medium components and culture conditions were identical to those described for regeneration assays. After three months of culture, adventitious shoot regeneration rate (%) was calculated as defined above.

### Agrobacterium-mediated transformation

*Agrobacterium tumefaciens* strain LBA4404 was used for genetic transformation. The binary vector pCAMBIA_KmR (Fig. S4a), carrying the *nptII* selectable marker, was used for transformation system establishment and optimization. To induce early flowering and thereby shorten the juvenile phase of intergeneric hybrids, the flowering-induction gene *MdFT1* ^56^ was overexpressed using the binary vector pPBV_Neo/Kana-CaMV35S-*MdFT1*-NosT (Fig. S4b), which harbors *nptII* and *MdFT1* driven by the CaMV 35S promoter.

Detailed transformation procedures are described in Supplementary Methods S1 according to Li et al. (2023) with minor modifications. To improve the recovery efficiency of regenerated shoots, Km concentrations of 5, 10, 15, 20 and 25 mg L^−1^ were evaluated at initial antibiotic selection after co-cultivation. Filter paper-assisted co-cultivation was subsequently tested at Km concentration of 5 mg L^−1^ to further enhance shoot recovery. To assess shoot recovery capacity under antibiotic selection and final transformation efficiency, two parameters were calculated: Regeneration rate (%) under antibiotic selection= (number of explants producing regenerated shoots / total number of inoculated explants) × 100; transformation efficiency (%) = (number of transgene-positive shoots/ total number of inoculated explants) × 100.

### Molecular confirmation and gene expression analysis

Genomic DNA was extracted from regenerated shoots of putative transformants, and the presence of the *nptII* and 35S-*MdFT1*-NosT transgenes was confirmed by PCR using primers (Supplementary Table S3). PCR was conducted in a 10 μL reaction volume containing 5 μL 2 × GoTaq Master Mixes (Promega, Tokyo, Japan), 100 nM each of the forward and reverse primers, and 10-50 ng of genomic DNA. The amplification program consisted of an initial denaturation at 94 °C for 3 min, followed by 35 cycles of 94 °C for 30 s, 58 °C for 30 s, and 72 °C for 40 s, with a final extension at 72 °C for 5 min. PCR products were analyzed by electrophoresis on 1.0% agarose gels. Wild type (WT) plants were included as negative controls. PCR-positive transgenic lines were maintained on shoot multiplication medium supplemented with 50 mg L^−1^ Km and 50 mg L^−1^ Me. Flowering was monitored under *in vitro* culture conditions, and the time to flower emergence was recorded relative to initial *Agrobacterium* infection.

For gene expression analysis, total RNA was extracted from transgenic lines and WT plants. Quantitative RT-PCR was carried out on a Takara Dice instrument (Takara, Shiga, Japan). The reaction mixture was prepared using TB Green Premix Ex Taq II. Primer sequences are listed in Table S3. Relative *MdFT1* expression levels were calculated using the 2^−^ΔΔCt method, with *MdACTIN* as the internal reference gene and WT plants as the calibrator.

## Supporting information

All the supplementary files

## Statistical analysis

Statistical analyses were performed using JASP software (version 0.95.4). Two-way ANOVA was applied to factorial experiments, whereas one-way ANOVA was used for single-factor analyses. When appropriate, Dunnett’s test (vs. control) or Tukey’s HSD test (pairwise comparisons) was applied. Differences were considered statistically significant at p < 0.05.

## Acknowledgements

The authors thank Drs. Sadao Komori and Masato Wada for valuable discussions. This research was supported by the Research and implementation promotion program through open innovation grants (JPJ011937) from the Project of the Bio-oriented Technology Research Advancement Institution (BRAIN) to TM and the Japan Society for the Promotion of Science JSPS [grant numbers 22K14887 to TM].

## Author contributions

F.L. and T.M. designed the experiments. T.M. constructed the vectors. F.L. performed the regeneration and transformation experiments. M.F. and M.F. performed the FT genetic transformation and phenotypic analyses. F.L. analyzed the data and drafted the manuscript. T.M., Y.W. and A.I. supervised the project and revised the manuscript.

## Data availability statement

All data generated or analyzed during this study are included in this present article.

