## Supplementary material for "A coordinated regeneration–selection strategy enables genetic transformation and rapid flowering in apple × pear intergeneric hybrids": All the supplementary files

**Supplementary Table S1. Effects of genotype and medium on callus induction rate (%)**

| Genotype | Medium | | | |
|---|---|---|---|---|
| | M1 | M2 | M3 | M4 |
| FjAD3-1 | 100.0 ± 0.0* a** | 98.7 ± 2.3 a | 98.0 ± 2.0 ab | 98.7 ± 2.3 a |
| FjAD3-2 | 91.7 ± 14.4 a-d | 97.5 ± 2.5 ab | 99.1 ± 1.4 a | 95.0 ± 6.6 a-c |
| FjYali B23-2 | 36.7 ± 10.0 g | 70.0 ± 14.5 c-f | 71.1 ± 10.2 b-f | 91.1 ± 10.2 a-d |
| FjOb3-1N | 98.9 ± 1.9 a | 99.2 ± 1.4 a | 95.6 ± 5.1 a-c | 93.3 ± 11.6 a-c |
| FjOb10-3 | 86.0 ± 5.3 a-e | 71.2 ± 4.7 b-f | 58.2 ± 16.4 fg | 84.8 ± 12.9 a-f |
| FjOb10-3_4N | 81.2 ± 8.0 a-f | 65.1 ±10.3 d-f | 61.2 ± 13.3 e-g | 95.4 ± 3.0 a-c |

| Two-way ANVOA | df | F | p |
|---|---|---|---|
| Genotype | 5 | 30.47 | < 0.001 |
| Medium | 3 | 7.49 | < 0.001 |
| Genotype × Medium | 15 | 6.09 | < 0.001 |

*Values are presented as mean ± SD.

**Different letters indicate significant differences among treatments ($p < 0.05$).

**Supplementary Table S2. Effects of genotype and medium on shoot regeneration rate (%)**

| Genotype | Medium | | | |
|---|---|---|---|---|
| | M1 | M2 | M3 | M4 |
| FjAD3-1 | 99.3 ± 1.2* a** | 65.3 ± 9.5 b-e | 95.3 ± 3.1 ab | 71.3 ± 13.6 a-d |
| FjAD3-2 | 79.2 ± 14.7 a-c | 79.2 ± 6.3 a-c | 86.7 ± 7.6 ab | 33.3 ± 10.4 e-i |
| FjYali B23-2 | 6.7 ± 6.7 i | 3.3 ± 3.3 i | 33.3 ± 17.6 e-i | 7.8 ± 5.1 i |
| FjOb3-1N | 73.9 ± 13.4 a-c | 53.6 ± 8.8 c-f | 35.0 ± 16.4 e-i | 53.1 ± 13.3 c-g |
| FjOb10-3 | 21.0 ± 6.6 hi | 16.3 ± 10.0 hi | 21.3 ± 2.3 g-i | 40.5 ± 16.5 d-h |
| FjOb10-3_4N | 25.2 ± 1.4 f-i | 15.3 ± 5.0 hi | 5.5 ± 5.7 i | 29.8 ± 14.2 f-i |

| Two-way ANVOA | df | F | p |
|---|---|---|---|
| Genotype | 5 | 98.16 | < 0.001 |
| Medium | 3 | 5.84 | 0.002 |
| Genotype × Medium | 15 | 7.89 | < 0.001 |

*Values are presented as mean ± SD.

**Different letters indicate significant differences among treatments ($p < 0.05$).

**Supplementary Table S3. Primer sequences used for PCR and RT-qPCR analysis**

| Gene | Primer | Sequence (5′–3′) | Product size (bp) | Application |
|---|---|---|---|---|
| *npt*II | nptII-F | ACAAGATGGATTGCACGCAGG | 780bp | PCR(Transgene detection) |
| | nptII-R | AACTCGTCAAGAAGGCGATAG | | |
| *FT*-TDNA | P35S-F | TGATGTGATATCTCCACTGAC | 799bp | PCR(Transgene confirmation) |
| | NosT-R | CTGTTGAATTACGTTAAGCATG | | |
| *MdFT1* | MdFT1-F | CCAAGTGACCCCAACCTAAA | 175bp | RT-qPCR(Gene expression analysis) |
| | MdFT1-R | CATACACYGTTTGCCTACCC | | |
| *ACTIN* | ACTIN-F | TGACCGAATGAGCAAGGAAATTACT | 222bp | RT-qPCR (reference gene) |
| | ACTIN-R | TACTCAGCTTTGGCAATCCACATC | | |

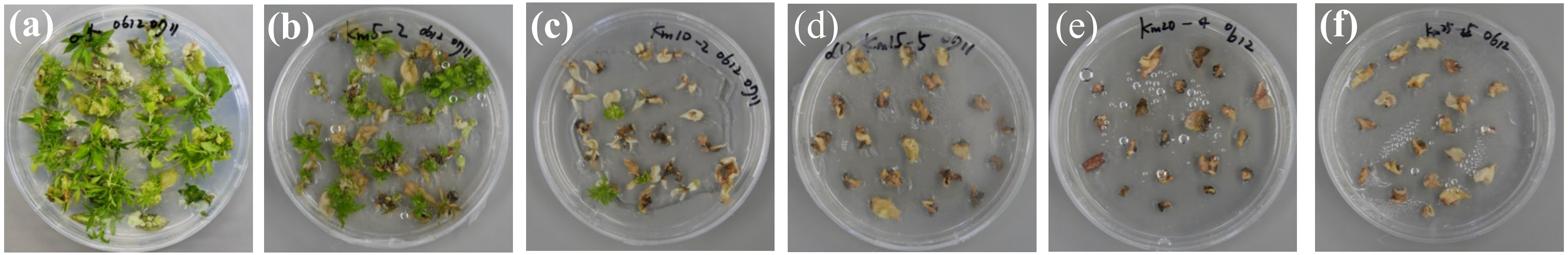


**Supplementary Fig. S1. Effects of increasing kanamycin concentrations on shoot regeneration in FjAD3-1**
(a)0 mg $L^{-1}$; (b) 5 mg $L^{-1}$; (c) 10 mg $L^{-1}$; (d) 15 mg $L^{-1}$; (e) 20 mg $L^{-1}$; (f) 25 mg $L^{-1}$.
Images were captured after three months of culture. Petri dishes were 90 mm in diameter.

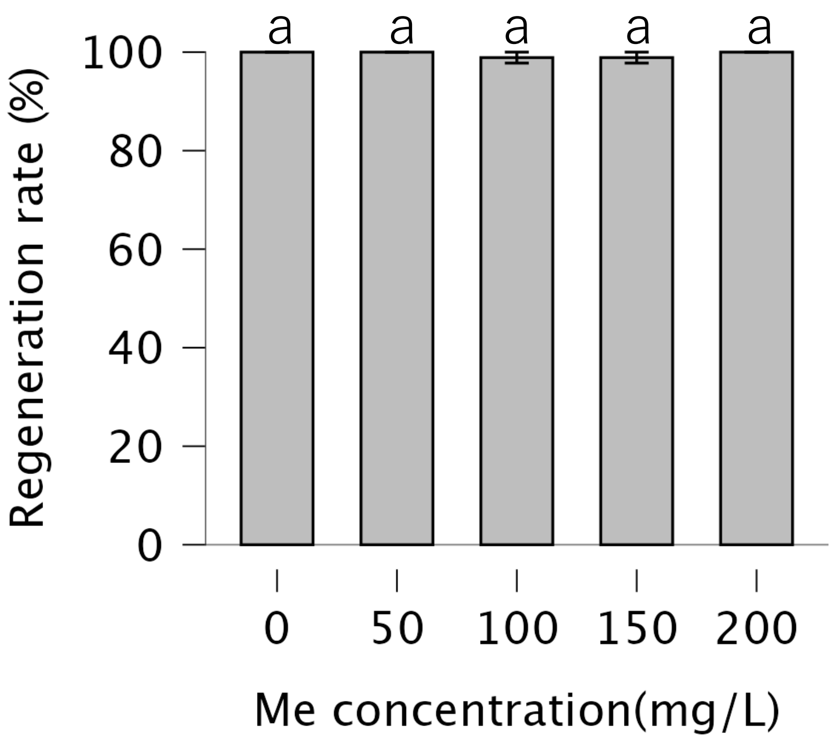


**Fig. S2. Shoot regeneration rates under different meropenem concentrations**
Data are presented as mean ± SD. Different letters indicate significant differences among treatments (Dunnett's test, $p < 0.05$).

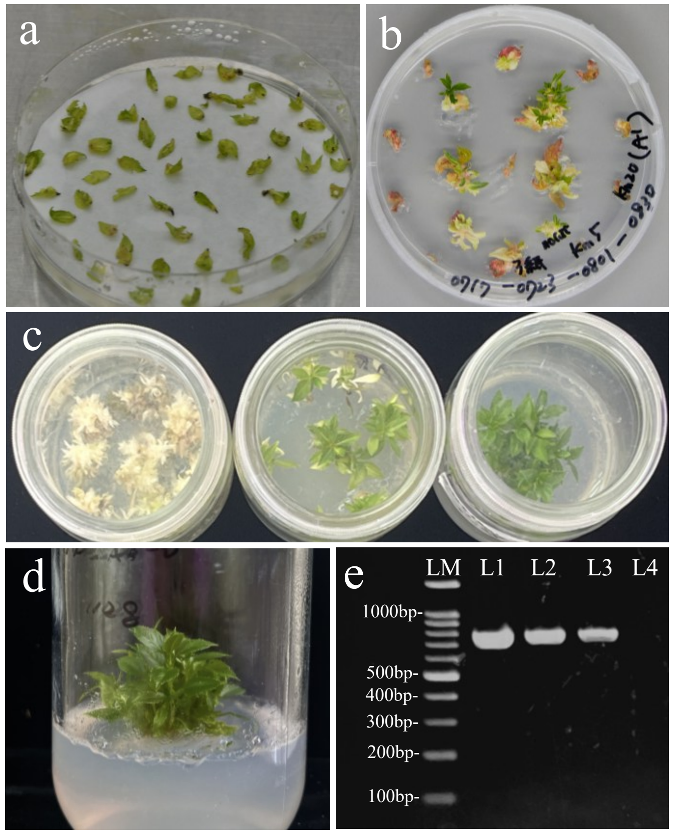


**Fig. S3. Stepwise selection and recovery of kanamycin-resistant shoots during Agrobacterium-mediated transformation**

(a) Co-cultivation of explants on regeneration medium(7days) with filter paper.

(b) Shoot regeneration under stepwise kanamycin selection (5 mg $L^{-1}$ followed by 20 mg $L^{-1}$).

(c) Multiplication of shoots on kanamycin-containing medium, showing bleached escape tissues (left), shoots under selection (middle), and green putative kanamycin-resistant shoots (right).

(d) Stable proliferation of putative transgenic shoots.

(e)PCR verification of the transgene in regenerated plants. Lane M: 100 bp DNA ladder (SMOBIO ExcelBand DM2100). Lane 1: plasmid containing the *npt*II gene used as a positive control. Lanes 2–3: regenerated plants showing the expected PCR amplification of the *npt*II gene (780bp). Lane 4: FjAD3-1 wild-type plant used as a negative control.

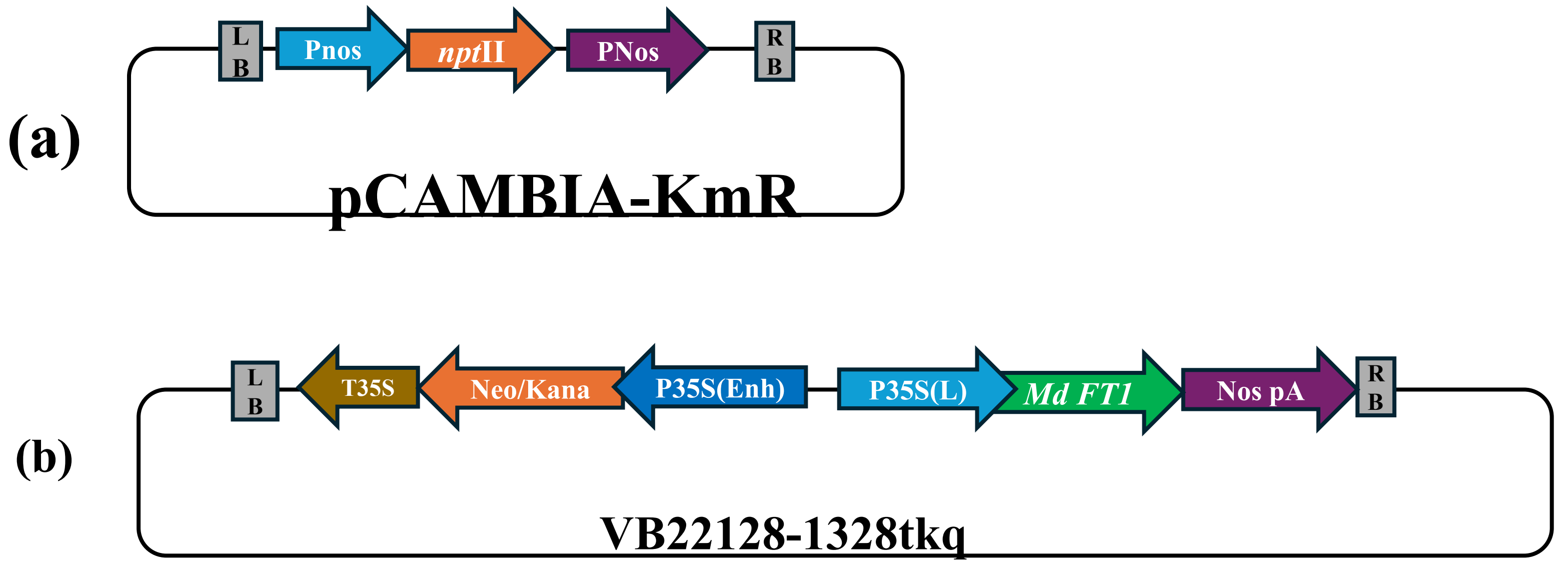


**Supplementary Fig. S4. Schematic diagrams of binary vectors used for Agrobacterium-mediated transformation**

(a) Vector pCAMBIA-KmR, carrying *npt*II gene, used during establishment of the transformation system.

(b) Vector pPBV-Neo/Kana-CaMV35S-MdFT1, carrying the early-flowering gene *MdFT1* and the *npt*II selectable marker.

**Supplementary Methods S1.**

Agrobacterium-mediated leaf transformation procedure:

Plasmids were introduced into *Agrobacterium tumefaciens* by heat shock (42°C for 90 s). A single colony harboring the target plasmid was cultured in liquid LB medium at 28°C with shaking at 200 rpm until $OD_{600}$ reached approximately 0.5. Bacterials were collected by centrifugation at 3000 rpm for 10 min, washed twice with liquid MS medium, and resuspended in liquid MS medium supplemented with 50 µM acetosyringone (AS). The suspension was adjusted to an $OD_{600}$ of 0.5 and incubated at 25°C with shaking at 100 rpm for 2 h prior to infection.

Young leaves from apple × pear intergeneric hybrids were gently wounded with a sterile blade and immersed in the bacterial suspension for 30 min. After infection, excess bacteria were removed using sterile filter paper, and explants were placed on regeneration medium M1 for co-cultivation. Co-cultivation was performed in the dark for 5–10 days and terminated when a thin bacterial film became visible on the medium surface.

Following co-cultivation, explants were incubated in liquid MS medium supplemented with 50 mg $L^{-1}$ meropenem for 30 min to eliminate residual *Agrobacterium*. Explants were then transferred to initial selection medium (M1 regeneration medium containing 5–20 mg $L^{-1}$ kanamycin and 50 mg $L^{-1}$ meropenem) and cultured in the dark. After 14 days, cultures were transferred to low-light conditions for 2 weeks. They were then moved to the final selection medium (M1 regeneration medium supplemented with 20 mg $L^{-1}$ kanamycin and 50 mg $L^{-1}$ meropenem) and maintained under a 16 h light / 8 h dark photoperiod. The final selection medium was renewed monthly.

Adventitious shoots exceeding 5 mm in length were transferred to shoot multiplication medium supplemented with 50 mg $L^{-1}$ kanamycin and 50 mg $L^{-1}$ meropenem. Shoots were subcultured monthly. All plant cultures were maintained at 25 °C.
